# Design of a genetic layered feedback controller in synthetic biological circuitry

**DOI:** 10.1101/647057

**Authors:** Chelsea Y. Hu, Richard M. Murray

## Abstract

Feedback control is the key to achieve robust performances for many engineered systems. However, its application in biological contexts is still largely unexplored. In this work, we designed, analyzed and simulated a layered controller functioning at both molecular and populational levels. First, we used a minimal model of three states to represent a system where state A activates state B; state R is a by-product of state B that acts as a negative feedback regulating both state A, B, and sequentially R. We call the feedback applied to state B a *cis* feedback and the one applied to state A a *trans* feedback. Through stability analysis via linearization at equilibrium and sensitivity analysis at transient state, we found that the *cis* feedback attenuates disturbances better but recovers slower; the *trans* feedback recovers faster but has more dramatic responses to fluctuations; the layered feedback demonstrates both advantageous traits of the two single layers. Then we designed two versions of synthetic genetic circuits to implement the layered controller in living cells. One version with an sRNA as regulator R, the other with a transcription factor protein as the regulator R. The analysis and dynamical simulation of the models confirmed the analytical results from the minimal model. At the same time, we found that the protein regulated feedback controls have faster recovery speed but the RNA version has a stronger disturbance attenuation effect.

## Introduction

One of the central goals of synthetic biology is to design and engineer biological systems with desired functions. Very much like other engineering disciplines, feedback control is useful as a mechanism to provide robust performance in these engineered biological systems. In nature, there is a wide range of biological phenomena that represent the successful utilization of feedback. For instance, bacteria move towards nutrients and away from toxic environments through chemotaxis signaling to ensure their survival [1]; the human body maintains a relatively constant glucose level in the bloodstream through insulin production by the pancreas; at a larger scale, predator-prey dynamics are the key to maintaining a stable ecosystem. These phenomena are all regulated by the principle of feedback: implement correcting actions based on the difference between desired and actual performance [2]. Feedback thus buffers systems from external disturbances and variations of components within the system. However, feedback can also destabilize the system when improperly designed [3]. Feedback design is especially challenging in biological systems due to its complexity — all molecular species are part of an extensive regulatory network that consists of numerous feedback mechanisms.

An accurate model is essential when designing feedback control in biological networks. However, a model that includes all the detailed molecule interactions in a living cell would be substantial and computationally expensive. In traditional engineering disciplines, reduced-order models are often used to outline the general dynamics of a system without describing every underlying mechanism [3]. Similarly, this approach of modeling is now widely used in synthetic biology to aid the design of simple biological circuits [4]. At the same time, our ability to build and test parts and modules of these circuits has been significantly enhanced over the past two decades. This is largely due to numerous advanced tools like next-generation sequencing, the discovery of fluorescent proteins [5] and modern cloning methods [6, 7, 8]. Consequently, the toolbox of characterized genetic components, modules, and motifs is expanding exponentially. As the complexity of synthetic circuits grows, its interplay with control theory becomes more prominent — proper design of feedback control is essential for a complex, sophisticated synthetic network, while the modularity and scalability of biological components offer valuable knowledge for the theoretical development of biological feedback control.

Just as nature deploys feedback controls, many well-designed engineering systems have multiple feedback controllers layered together to function as a system. However, the design principle of layered feedback in biology is mostly unexplored. In this work, we investigated the theoretical properties of a *trans-cis* layered feedback controller in living cells. Starting with a minimal model, we used a common control theory approach to analyze the stability of *cis*, *trans* and layered feedback controllers. We found that the *cis* feedback trades off speed for stability, while the *trans* feedback trades off stability for speed. When the two types of feedback are layered together, the controller shows an integrated behavior that accentuates both advantages of the two individual layers. We then designed two synthetic circuits that would carry out this design at the molecular level in living cells, with an RNA-based regulator and a protein-based regulator. The result showed that the RNA version has a stability advantage while the protein version has a recovery-speed advantage. These results serve as experimental guidance for the next stage where we will build and test these designs. We hope that the models and experimental results together will help us understand and predict the behavior of layered controllers in synthetic biology.

## Results and Discussion

### Stability analysis on a layered feedback with a minimal model

In order to investigate the cooperative behavior of two layers of negative feedbacks, we started our analysis with a minimal model of three species. As shown in Figure 1A, we defined two nodes A and B, where B is the observable output of the system that is activated by A. R is a by-product of species B that negatively regulates expression of B and itself through two possible routes: *cis* feedback (R represses B) and *trans* feedback (R represses A). The activation and repression here are modeled with first order Hill functions. Both A and B are assumed to be protein species while R could be either a protein species or a regulator RNA species. Our analysis in the rest of this work focuses on four designs: no feedback, *cis* feedback only, *trans* feedback only and layered feedback.

**Figure 1:**
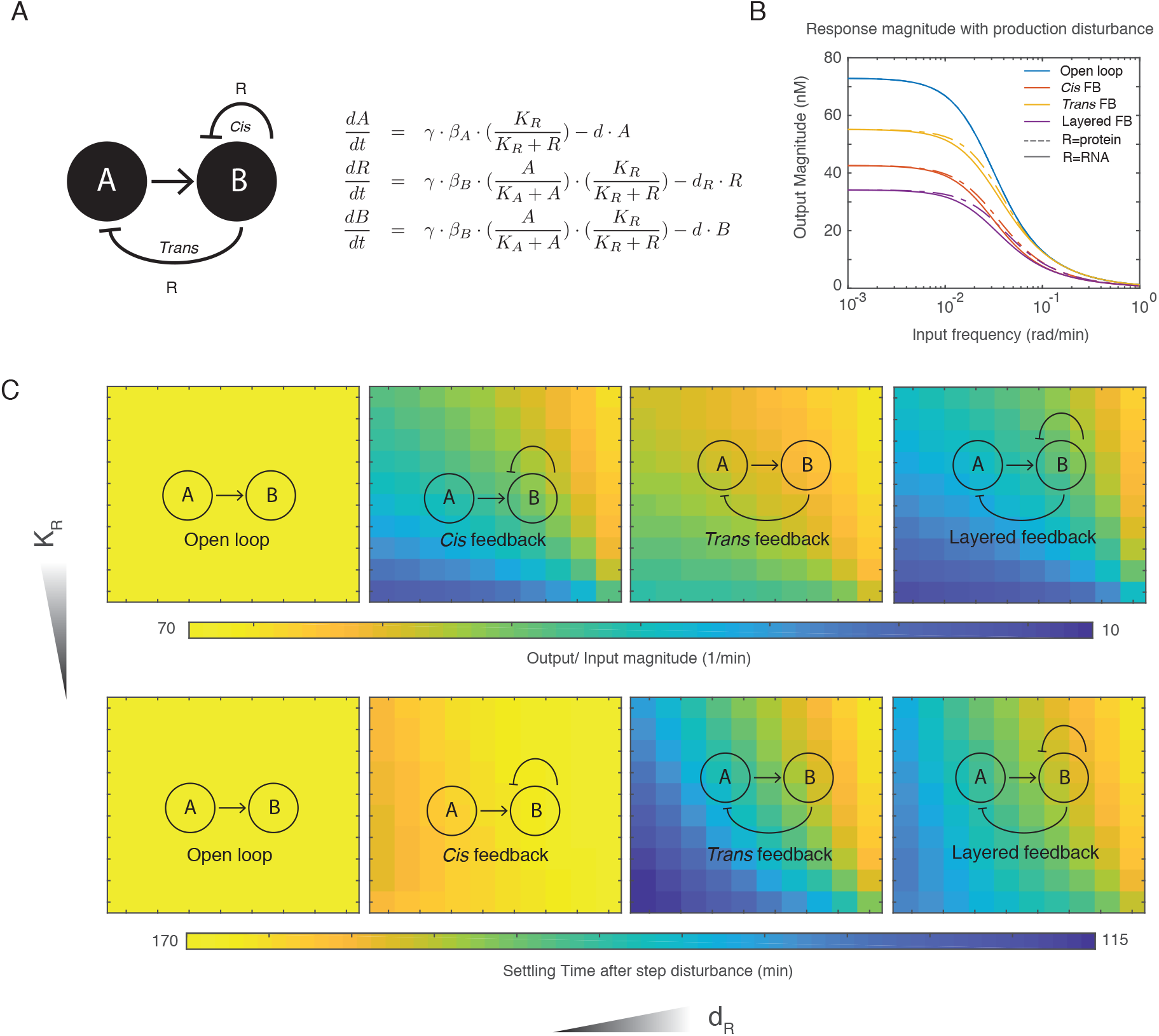
Stability analysis of the layered feedback controller using a minimal model. (A) Schematics and ODEs of the system described by a minimal model. Here A and B are two molecular species, where A promotes the expression of B. R is a by-product of B that acts as a regulator. R negatively regulates species B through *cis* feedback and regulates species A through *trans* feedback. When both types of feedback exist in the same system, it is termed layered feedback. (B) Response magnitude with production disturbance at different frequencies. This plot shows the magnitude response of species B (y-axis) when all three species’ production rates are subjected to a perturbation at different frequencies (x-axis). The blue, yellow, red, and purple curves represent open loop, *cis* feedback, *trans* feedback, and layered feedback circuit structures, respectively. Solid lines represent the magnitude response of these four structures when regulator R is an RNA species, while dashed lines represent protein species. (C) Disturbance attenuation and settling time of four circuit constructs in a two-dimensional parameter space *K*_*R*_ and *d*_*R*_. Top four panels show the output disturbance magnitude when all three species’ production rates are subjected to a perturbation at low frequency. Bottom four panels show the time it takes for the dynamic to settle back to equilibrium after the disturbance. Both response magnitude and settling time are represented by the color schemes (see color bar). For each of the eight heat maps, the x-axis (*d*_*R*_) represents the degradation rate of regulator R; a large *d*_*R*_ represents a regulator with fast degradation rate. The y-axis (*K*_*R*_) represents the repression constant of regulator R; a large *K*_*R*_ represents a regulator with weak repression strength.

First, we performed a stability analysis of these four constructs at equilibrium using their transfer functions. The transfer function describes the relation between inputs and outputs of a linear system in the frequency domain [3]. Since all of the four systems are nonlinear, we linearized them at their equilibrium points before converting to the frequency domain. Here we define the output as species B. The input is defined as *γ*, which is a unitless scalar that impacts the expression rate of all three species (*β*_*A*_ and *β*_*B*_). In Figure 2B, we show the magnitude response of these four systems in the frequency domain. In this figure, the x-axis represents the input frequency in rad/min, and the y-axis represents the output/input magnitude, which has a unit of nM. When the input frequency is low, this could be understood as a long-lasting change to the universal expression rates. When the input frequency is high, the input disturbance oscillates rapidly, and output response to the disturbances diminishes. In this figure, we show that the *cis* feedback (red) attenuates disturbances better than the *trans* feedback (yellow). The layered feedback shows the strongest attenuation effect (purple). We also observed that there is only a slight difference between an sRNA regulator (solid lines, with *K*_*R*_ = 10, *d*_*R*_ = 0.3) and a protein regulator (dashed lines, with *K*_*R*_ = 100, *d*_*R*_ = 0.03) at intermediate input frequencies.

**Figure 2:**
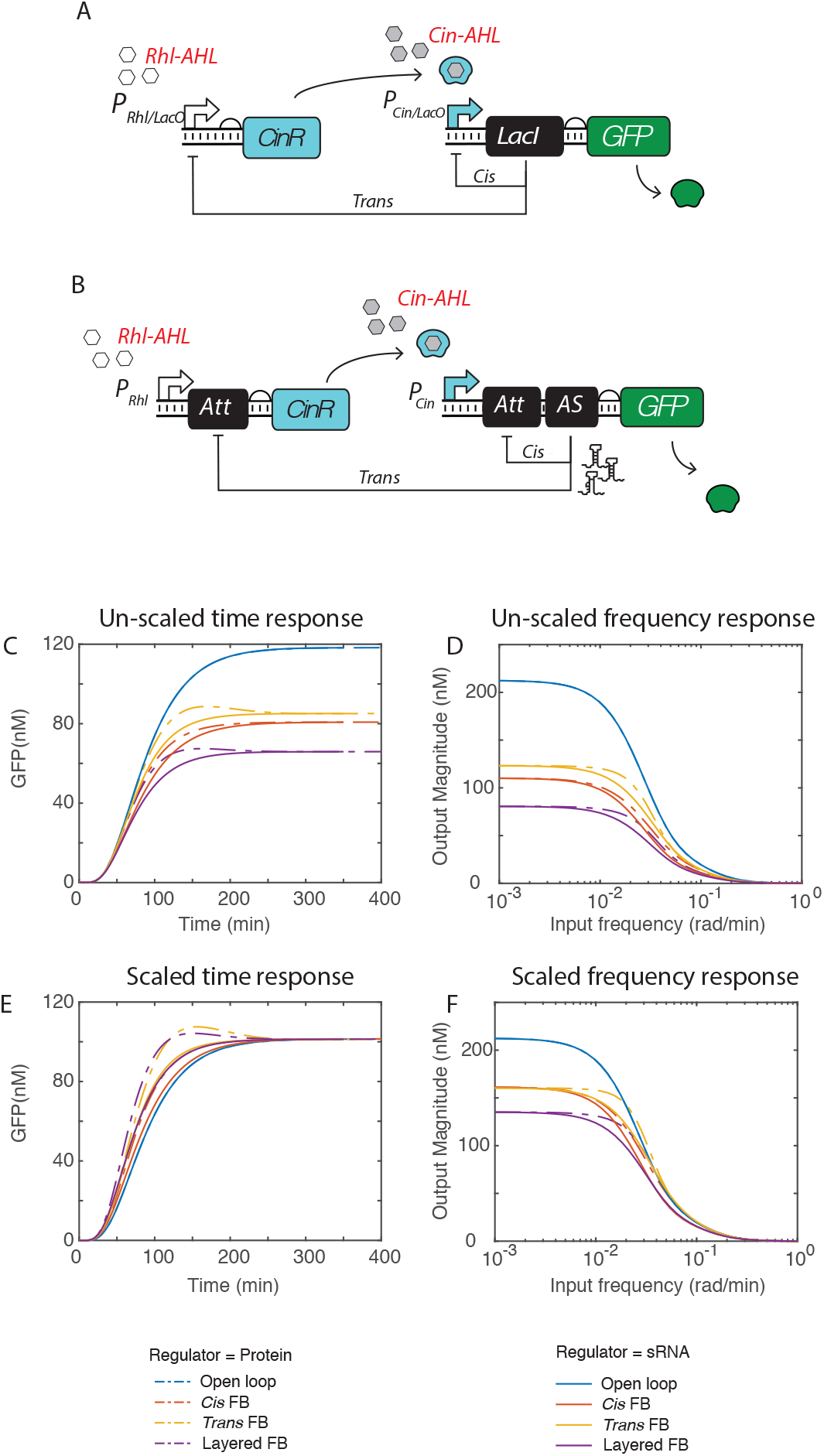
Design and RBS scaling for molecular layered feedback. (A) Schematics of the layered feedback controller at the molecular level, with a protein regulator. To compare this design with the minimal schematics in Figure 1A, CinR here is species A, GFP is species B, and LacI is regulator R. LacI regulates the transcription of CinR and GFP through its interaction with promoter P_Rhl/LacO_ and promoter P_Cin/LacO_. (B) Schematics of the layered feedback controller at the molecular level, with an sRNA regulator. Species R is an sRNA regulator that interacts with the attenuator (Att) RNA. This interaction causes transcriptional termination, and therefore down regulates the gene expression of downstream genes. (C)-(D) Dynamic of GFP expression (C, E) and response magnitudes (D, F) of open loop, *cis* feedback, *trans* feedback and layered feedback circuit structures (four colors) with both protein and RNA regulators (dashed and solid lines). (C) and (D) are the dynamics and magnitude response without RBS scaling; each circuit structure has a unique equilibrium. (E) and (F) are the dynamics and magnitude response with RBS scaling; all four constructs have the same equilibrium.

In Figure 1C, we plotted the magnitude response of each construct at low frequency in a two dimensional parameter space *K*_*R*_ (y-axis) and *d*_*R*_ (x-axis). We found that over a wide parameter space, the *cis* feedback attenuates disturbances at low frequency better than the *trans* feedback, while the layered feedback shows the most attenuation. In each feedback design, regulators with strong repressive strength and a slow degradation rate achieve the most disturbance attenuation. In the lower panel of Figure 1C, we plotted the settling time of each construct after step disturbance in the same two dimensional parameter space *K*_*R*_ (y-axis) and *d*_*R*_ (x-axis). We found that although the *cis* feedback has strong attenuation of disturbance, it takes longer to recover back to equilibrium, while the *trans* feedback recovers back to equilibrium faster but attenuates the disturbances less effectively. The layered feedback, on the other hand, appears to have a moderate settling time that is in between the *cis* and *trans* feedback mechanisms.

### Layered feedback design at the biomolecular level

We designed two versions of the layered feedback controller, one with a transcription factor protein regulator (Figure 2A), the other with an sRNA regulator (Figure 2B). As shown in Figure 2A, the system is induced with both Cin-AHL and Rhl-AHL. The transcription of CinR is controlled by promoter P_Rhl/LacO_, and the transcription of GFP and LacI is controlled by promoter P_Cin/LacO_ [9]. Compared with the minimal design in Figure 1A, here CinR is species A, GFP is species B and LacI is species R. We assume that RhlR is encoded in the genome and constitutively expressed. In the RNA regulator design (Figure 2B), the system is also induced with both Cin-AHL and Rhl-AHL. The transcription of CinR is controlled by promoter P_Rhl_, and the transcription of GFP and LacI is controlled by promoter P_Cin_. A piece of attenuator RNA is placed upstream of both CinR and GFP, and an sRNA repressor AS is placed in between GFP and the attenuator as the regulator. When AS is present, it interacts with the attenuator RNA (Att) to cause a conformation change that terminates transcription of the downstream genes. In this design, AS is the regulator species R in Figure 1A.

We first simulated and compared the dynamics of each of the two controllers separately. As shown in Figure 2C, we simulated four designs: open loop (blue), *cis* feedback (red), *trans* feedback (yellow) and the layered feedback (purple) for both of the designs (solid lines for RNA regulator, dashed line for protein regulator). As expected, different feedback designs have different equilibrium points. We performed stability analysis with transfer functions on these constructs as we did in Figure 1, and the result is similar to that of the minimal model. However, the variation in equilibrium of these four systems could be problematic for two reasons: (1) when the output protein B in Figure 1A has a downstream impact (e.g. metabolism, regulation or cell fate, etc.), species might have an optimal equilibrium to ensure the functionality of the system. Deviating from this optimal equilibrium might impact the broad system’s overall behavior. (2) Because these four constructs have different equilibrium points, it is difficult to conclude what is causing the disturbance attenuation. As shown in Figure 2D, layered feedback has the strongest disturbance attenuation effect, but it also has the lowest equilibrium (Figure 2C). It is not clear whether the disturbance attenuation is caused by the feedback control topology or by the lower equilibrium point. Fortunately, protein expression is determined by both transcription and translation in biology. This allows us to adjust the equilibrium levels of the four constructs in the translational step. As shown in Figure 2E, we matched the equilibrium states of CinR and GFP across all four constructs by re-scaling the RBS strengths for these two proteins. The RBS scaling factors are listed in Table 4. Note that the table represents the scaling factors used for both protein and RNA versions of the four constructs. This is due to the intentional parameter choices for the model, which assumes both protein and RNA regulators achieve the same repressive strength, despite the difference in degradation and maturation rates. This scaling step allowed a fair comparison of the four topologies as the feedback’s impacts on expression strength of CinR and GFP were removed.

**Table 1:** Reduced model parameters

| Parameters | Description | Unit | Guess |
| --- | --- | --- | --- |
| $\beta_A$ | production rate of A | nM/min | 1 |
| $K_R$ | repression constant of R | nM | 10/100 |
| $d$ | degradation rate governed by cell division | 1/min | 0.03 |
| $\beta_B$ | production rate of both R and B | nM/min | 5 |
| $K_A$ | activation constant of A | 1/min | 100 |
| $d_R$ | degradation rate of R, always greater than or equals to $d$ | 1/min | 0.3/0.03 |
| $\gamma$ | universal production scaling factor | NA | 1 |

**Table 2:** Molecular level layered control model species

| Species | Description |
| --- | --- |
| $M_s$ | mRNA of signaling protein (CinR) |
| $S$ | signaling protein translated peptides (CinR) |
| $A$ | folded CinR protein bounded with AHL |
| $R$ | antisense RNA repressor |
| $M_G$ | GFP mRNA |
| $G$ | translated GFP peptides |
| $G_m$ | mature GFP (fluorescent) |
| $R_m$ | regulator mRNA |
| $R_p$ | regulator protein |

**Table 3:**
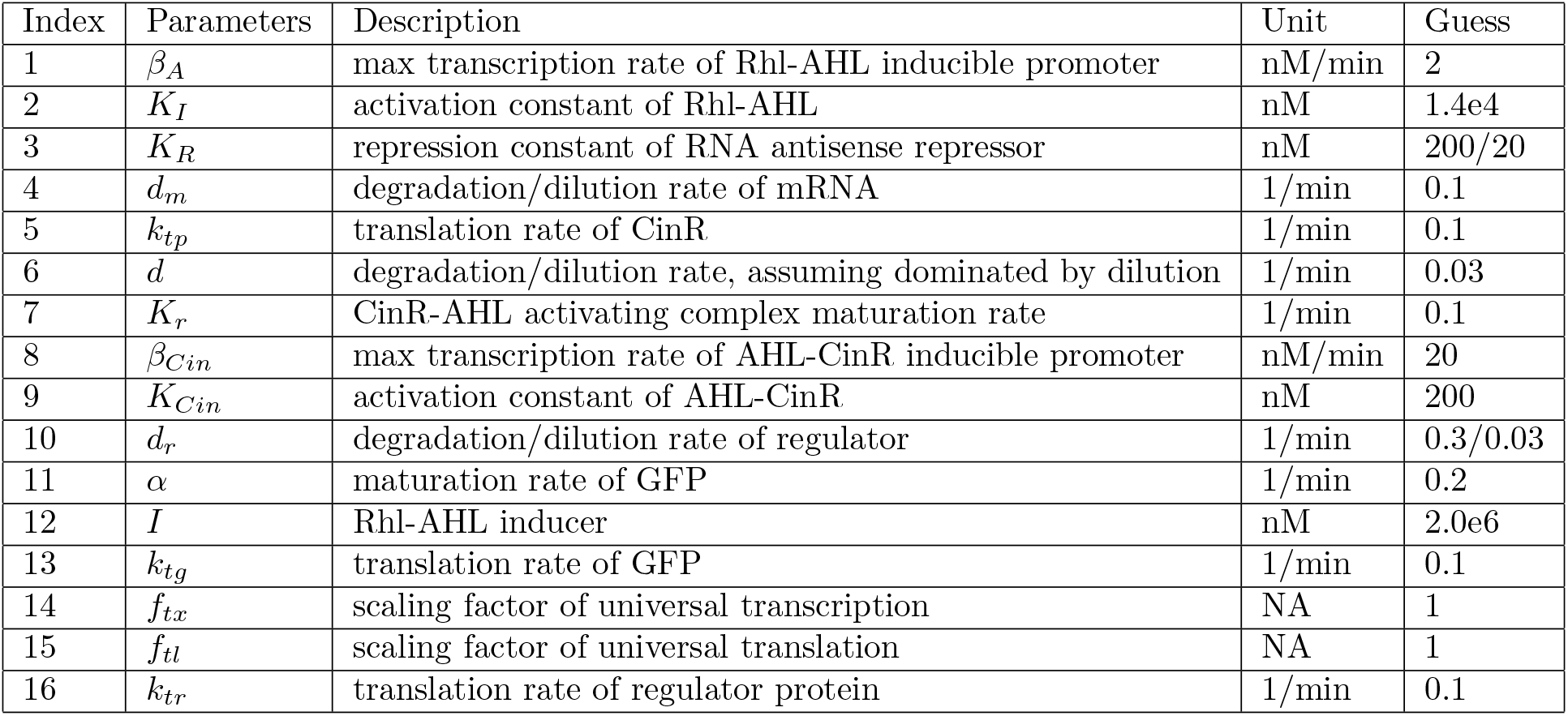
Model parameters

**Table 4:** RBS scaling

| RBS scaling factors | open loop | *cis* feedback | *trans* feedback | layered feedback |
| --- | --- | --- | --- | --- |
| CinR | 1 | 1 | 1.6875 | 1.4688 |
| GFP | 1 | 1.4648 | 0.9961 | 1.4609 |

In Figure 2E, we matched the equilibrium of each feedback construct to the open loop case. We observed that all three types of feedback speed up the response of the circuit, with *trans* feedback being the fastest. Additionally, the protein regulator feedback has a faster response time than those based on sRNA regulators. This result was observed from both the scaled (Figure 2E) and un-scaled circuit dynamics (Figure 2C), demonstrating that the fast dynamics with feedback are not entirely due to lower equilibrium values. We also noticed that as we performed that same stability analysis on RBS-scaled designs (Figure 2F), the response magnitudes of *trans* and *cis* feedback overlapped at low disturbance frequency. However, the disturbance attenuation effect, although less prominent in comparison to the un-scaled case, is still significant. The layered feedback also has a stronger disturbance attenuation at low input frequency than the two single feedbacks for both the scaled and un-scaled cases. This result indicates that the advantage of disturbance attenuation *cis* feedback has over *trans* feedback is likely due to its lower equilibrium, while the advantage of disturbance attenuation of layered feedback over single feedback is due to its system architecture.

This analysis also showed the key difference between RNA regulator mediated feedback and protein regulator mediated feedback. As presented in Figure 2C and 2E, feedback designs with protein-based regulators have a faster rise than the ones with RNA-based regulators. Based on the magnitude analysis results in Figures 2D and 2F, the two sets of designs do not show a significant difference at the lowest and the highest ends of disturbance frequency. In the middle range, RNA regulators seems to provide a slight advantage of disturbance attenuation.

Next we performed parameter sensitivity analysis [10, 11, 12] of the two types of layered feedback controllers in their transient states—with sRNA regulators and protein regulators. The sensitivity of parameters describes how much impact each parameter has to the observable state at a given time point. In other words, how much the observable state changes when each parameter is being perturbed. In Figures 3A-C, each heat map represents a sensitivity matrix of a certain construct. The x-axis represents time and the y-axis represents parameter index. Parameters are listed in Table 3. Parameters that are not involved in a given construct are greyed out. The color scheme represents the sensitivity of parameters to the output GFP. We found that the protein-based layered feedback controller increased the sensitivity of *k*_*tg*_ and *f*_*t*_*l* (Figure 3B), which indicates that it is less stable against translational fluctuation during the transient state. For both open loop and protein based layered feedback, this translational impact is long-lasting. By comparison, sRNA based layered feedback has a very different sensitivity profile (Figure 3C). It is significantly less dependent on translational parameters (*k*_*tg*_,*f*_*tl*_), and is more sensitive to a handful of transcriptional parameters (*d*_*m*_,*β*_*lux*_,*K*_*lux*_,*f*_*tx*_). However, the sensitivity of these parameters decreases rapidly over time as the circuit dynamic approaches equilibrium.

**Figure 3:**
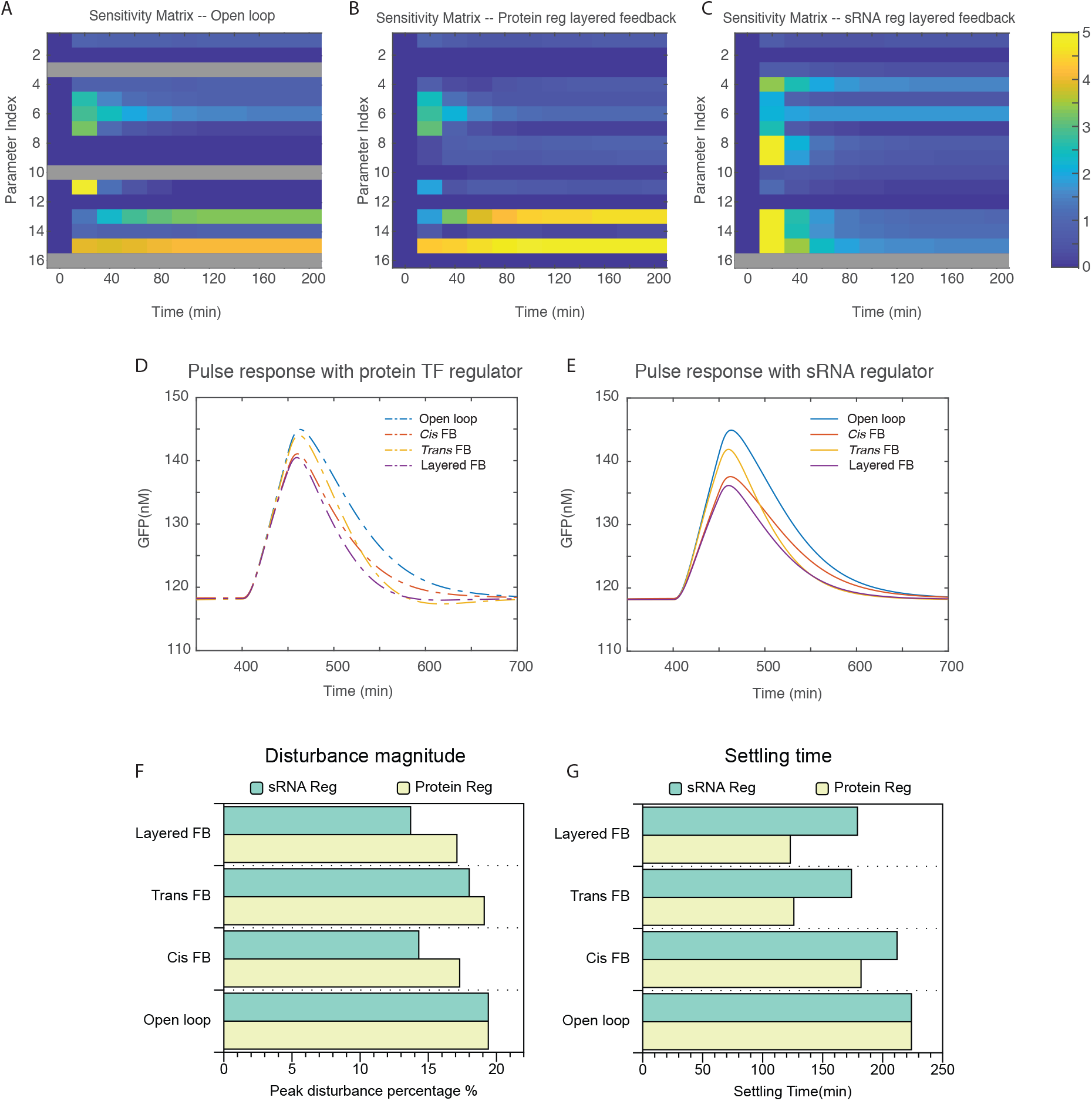
Sensitivity of parameters at system transient state and circuit response with a disturbance pulse. (A)-(C) Comparison of parameter sensitivity of open loop circuit (A) with two versions of layered feedback circuits: (B) protein regulators and (C) sRNA regulators in transient state. The y-axis represents the index of parameters that are listed in Table 3, x-axis is time. (D) Dynamics of the four constructs with protein regulators in response to a pulse of transcriptional disturbance. (E) Dynamics of the four constructs with sRNA regulators in response to a pulse of transcriptional disturbance. (F) Measured disturbance response magnitude of the four structures with sRNA regulators (green bars) and protein regulators (yellow bars), measured from simulation shown in (D) and (E). This figure shows that *cis* feedback attenuates disturbances better than *trans* feedback. Also, sRNA regulators in all three feedback designs show a slight advantage in disturbance attenuation. (G) Measured settling time of the four structures with sRNA regulators (green bars) and protein regulators (yellow bars), measured from the simulation shown in (D) and (E). This figure shows that *cis* feedback recovers slower than *trans* feedback after disturbances. Furthermore, protein regulators in all three feedback designs show a slight advantage in recovery speed.

Finally, we looked at the dynamic of feedback controllers in response to a transcriptional disturbance pulse via simulation (Figures 3E-G). We first scaled the RBS for all four constructs to match their equilibrium points. At equilibrium, we gave the universal transcriptional rate *f*_*tx*_ a 25% increase and held it for 50 minutes (Figures 3D,E). Then we plotted each construct’s percentage GFP response at its peak (Figure 3F) and the settling time after the disturbance (Figure 3G). Similar to the result we obtained from the minimal model in Figure 1C, we found that the *cis* feedback is less prone to disturbances but has longer settling time than *trans* feedback. When these two feedback controllers are layered together, the system inherits the strong disturbance attenuation from *cis* feedback and the fast recovery speed from *trans* feedback. Meanwhile, we found that the protein based feedback exhibits faster recovery but are less effective at noise attenuation than RNA based feedback. This observation is consistent with the analysis we presented in Figure 2.

Ordinary differential equations for sRNA mediated layered feedback controller:

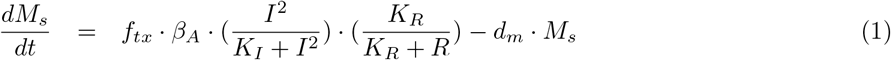

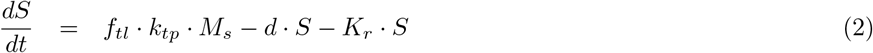

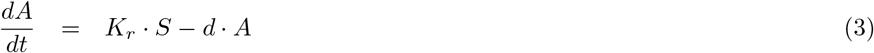

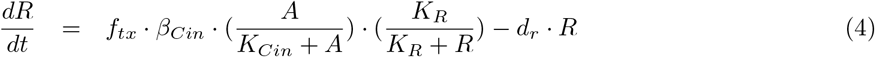

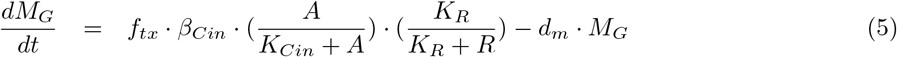

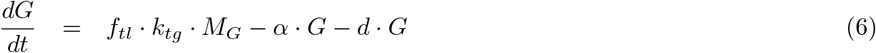

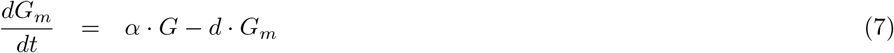

Ordinary differential equations for transcription factor protein mediated layered feedback controller:

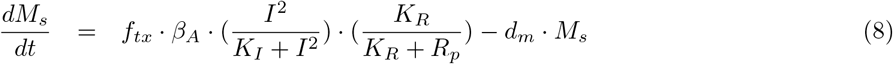

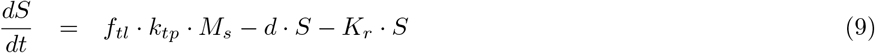

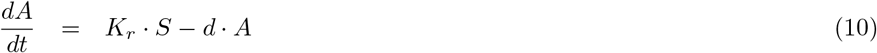

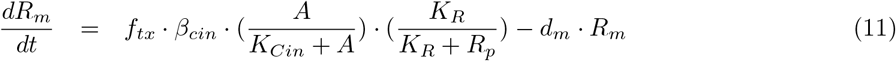

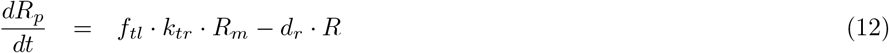

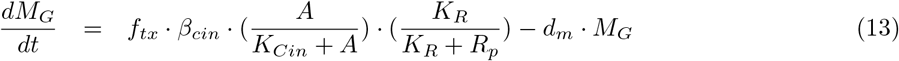

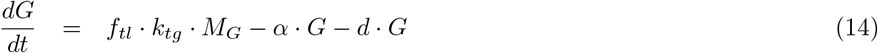

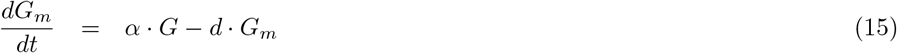

## Conclusion

With a control theory approach, we analyzed the properties of different types of negative feedback controls and the cooperative behavior shown when they are layered together. We concluded that the *cis* feedback compromises recovery speed for a stronger disturbance attenuation while the *trans* feedback sacrifices attenuation strength for faster recovery speed. As they are layered together, the layered feedback acquires a new property that demonstrates a combination of strong disturbance attenuation and fast recovery. This analysis was performed on both a minimal model and a reduced molecular model.

Like many reduced models, our model was based on numerous assumptions. Some of these assumptions help to reduce the model and offer valuable insights to the system. Others might fail to capture the essence of the system under certain conditions. To capture the accurate behaviors of controllers in biology, it is important to consider the molecular, even cellular environments of the system. This work is still in its early stages of development. In the next phase of this project, we will expand the model to include natural environmental feedback including resource constraints, cell division, and cell death. We anticipate that the designs should have very different behaviors when they are operating near resource capacities. We plan to explore strategies to preserve desired functionality of the controllers in the dynamical molecular environment of living cells. Finally, we plan to implement the design in cells to experimentally test these theoretical findings. We envision that the model would provide insightful guidelines for constructing the controllers, while the experimental results would offer a profound perception of the behavior of layered feedback controllers in biology.

## Acknoledgement

The authors would like to thank Reed McCardell, Ayush Pandey and Xinying Ren for their insightful discussions. The author C. Y. Hu is supported by Defense Advanced Research Projects Agency (Agreement HR0011-17-2-0008). The content of the information does not necessarily reflect the position or the policy of the government, and no official endorsement should be inferred.

